# Metabolomics reveals nucleoside analogs for regulating mucosal-associated invariant T cell responses

**DOI:** 10.1101/2023.01.30.526332

**Authors:** Shouxiong Huang, Manju Sharma, Larry Sallans, Chunshun Li, Zaman Kh, Divaker Choubey, David Newburg, Moises A. Huaman, Ted Hansen, Shugeng Cao

## Abstract

Although mucosal-associated invariant T (MAIT) cells recognize riboflavin-like metabolites from Gram-negative bacteria, MAIT cell stimulation by broad bacterial families and mammalian cells suggests the existence of novel ligands from different biological sources. Here we established a comparative platform of functional metabolomics and used *Mycobacterium tuberculosis* as a model to characterize novel metabolites for MAIT cell activation. We extracted and fractionated small metabolites of *M. tuberculosis* using high-performance liquid chromatography, showing a different MAIT cell stimulation pattern of *M. tuberculosis* metabolite fractions in comparison with *Escherichia coli* fractions. Mass profiling predicted multiple nucleoside analogs enriched in a biologically active fraction of *M. tuberculosis*. Whereas the synthetic forms of these predicted *M. tuberculosis* nucleosides were unavailable, structural-based autodocking of analogous nucleosides conserved in mammals supported potential binding with MR1 protein. Indeed, functional assays of these conserved nucleosides demonstrated guanosine as a stimulator and deoxyformyluridine as an inhibitor of MAIT cell activation. Identification of bioactive nucleoside metabolites broadly conserved in bacterial and mammalian systems will facilitate an understanding of the regulatory roles of MAIT cells in infectious and inflammatory conditions.

## Introduction

Conventional T cells recognize the peptide antigens presented by major histocompatibility complex (MHC) molecules to combat infections and regulate inflammation. In primary infections, conventional T cells require prolonged clonal cell expansion to generate sufficient numbers of pathogen-specific T cells to achieve effective immune responses against pathogens (Hansen et al., 2007; Roche and Cresswell, 2016). Unlike conventional T cells, mucosal-associated invariant T (MAIT) cells recognize non-peptidic metabolites presented by monomorphic MHC class I-related protein 1 (MR1) in response to bacterial pathogens and tissue inflammation (Huang, 2016; Huang et al., 2009; Kjer-Nielsen et al., 2012; Le Bourhis et al., 2010; Magalhaes et al., 2015). Like innate immune cells, MAIT cells exist in high numbers and express conserved activation receptors that allow a rapid response to various bacterial pathogens in primary infections (Le Bourhis et al., 2010; Reantragoon et al., 2013; Sharma et al., 2020). This rapid MAIT cell response at the early stage of bacterial infections or MR1 ligand stimulation protects the host against infections or regulates inflammations (Huang, 2016; Le Bourhis et al., 2010; Sakai et al., 2021). Beyond bacterial infections, various inflammatory conditions, autoimmune diseases, and cancers are also associated with the alteration of MAIT cell frequency and response (Chiba et al., 2018; Godfrey et al., 2019; Magalhaes et al., 2015). However, how MAIT cells contribute to immune responses in non-infectious diseases is unknown, partially due to the uncharacterized MAIT cell stimulants of mammals. Thus, identifying the novel and conserved stimuli for MAIT cells will facilitate elucidating the roles and mechanisms of MAIT cell responses in various infectious, autoimmune, and cancer diseases.

Currently, protective roles of MAIT cells have been demonstrated in bacterial lung infections in mice, including legionellosis (Wang et al., 2018), tularemia (Zhao et al., 2021), sepsis (Trivedi et al., 2020), as well as mycobacterial infections (Le Bourhis et al., 2010; Sakai et al., 2021). MAIT cells can elicit antimicrobial responses against gut bacterial infections as well, including *E.coli, Shigella flexneri* (Kurioka et al., 2015; Le Bourhis et al., 2010), and *Salmonella typhimurium* (Chen et al., 2017). For mycobacteria, MAIT cells protect mice by inhibiting the growth of nontuberculous *M. abscessus* (Le Bourhis et al., 2010; Martins et al., 1999) and *M. bovis* (Chua et al., 2012; Sakala et al., 2015), and partially suppress *M. tuberculosis* infection in mice (Sakai et al., 2021). These recent studies highlighted MAIT cells as promising targets to induce immune protection against various respiratory and intestinal bacteria by recognizing and responding to antigenic compounds that are expressed by different genera of bacteria. However, it is unknown whether MAIT cells only recognize bacterial antigens or whether MAIT cells can respond to both bacterial and mammalian stimuli. In uninfected immunopathological conditions, MR1-dependent antigen presentation is also critical for MAIT cells to produce interleukin (IL)-2 (Huang et al., 2005; Huang et al., 2008; Le Bourhis et al., 2010), IL-17a, and interferon γ (IFNγ) (Toubal et al., 2020), respond to an agonist (Chiba et al., 2017), and regulate MAIT cell responses in cancer (Gherardin et al., 2018; Petley et al., 2021). The identities of endogenous and novel bacterial regulators for MAIT cell activation remain to be elucidated for understanding MAIT cell functions in human diseases.

Different from peptide antigens for conventional T cells or lipid antigens for CD1-restricted T cells, multiple water-soluble metabolites have recently been identified as MAIT cell antigens. The bacterial antigens known so far for MAIT cells are intermediate metabolites in the riboflavin metabolic pathways, including ribityluracil (5-A-RU, 5-Amino-6-D-ribitylaminouracil) identified from multiple bacteria, including *E. coli* (Corbett et al., 2014), ribityllumazine identified from *Salmonella* and *E. coli* (Corbett et al., 2014; Kjer-Nielsen et al., 2012), and photo-lumazine from *Mycobacterium smegmatis* (Harriff et al., 2018). Abundance of these metabolites remains to be determined from various bacterial species, which express different protein sequences and levels of various riboflavin metabolic enzymes (Mortl et al., 1996; Wang et al., 2015; Zylberman et al., 2006), potentially leading to differential production of intermediate metabolites in the riboflavin pathway. Beyond these vitamin B metabolites (Kjer-Nielsen et al., 2012), multiple drug or drug-like compounds further expand the chemical diversity of MAIT cell antigens from natural lumazine or uracil compounds to synthetic aromatic and ketone compounds (Keller et al., 2017). However, how chemically diverse MAIT cell antigens are translated to MAIT cell responses in physiological or diseased conditions is unknown.

MAIT cells were initially thought to respond to a limited number of ligands, mainly because of the invariant TCRα chain that conveys the name of invariant T cells (Porcelli et al., 1993; Treiner et al., 2003). This notion seems reasonable while considering highly conserved innate receptors such as Toll-like receptors that bind a limited number of pathogen-associated molecular patterns (PAMPs) (Janeway, 1989). However, another possibility is that MAIT cells respond to various metabolite antigens, as similar to that natural killer T cell response to a variety of lipid ligands (Rossjohn et al., 2015). Indeed, despite a semi-invariant TCRα chain, MAIT cells express various TCRβ chains in response to *M. smegmatis, S. typhimurium*, and *C. albicans* with riboflavin pathways (Gold et al., 2014), as well as *Streptococcus pyogenes* without riboflavin biosynthetic enzymes (Meermeier et al., 2016). This evidence together with the identified drug-like MR1 ligands (Keller et al., 2017) and the unidentified endogenous MR1 ligands (Huang et al., 2005; Huang et al., 2009) suggests the presence of novel bacterial and mammal metabolite antigens other than riboflavin precursor metabolites, which we sought experimentally in this report.

We established a comparative metabolomics platform by integrating bacterial metabolite extraction, functional stimulation, mass profiling, ligand docking, and structural analyses to discover nucleoside compounds or analogs for regulating MAIT cell responses. We started by distinguishing the chemical composition of *M. tuberculosis* stimulants versus *E.coli* stimulants for MAIT cell activation. Elution and fractionation of small metabolites from *M. tuberculosis* using size exclusion and high-performance liquid chromatography (HPLC) provided *M. tuberculosis* eluates and unique fractions that activated MAIT cells. Mass spectrometry predicted the enriched nucleoside compounds from an active *M. tuberculosis* fraction. However, the predicted *M. tuberculosis* nucleosides are unavailable, and we sought to test the structurally similar nucleoside analogs, which are conserved in bacterial and mammalian cells. Indeed, functional assays showed that guanosine activated and deoxyformyluridine inhibited MAIT cell responses, respectively. These results expand the natural regulators of MAIT cell response from bacterial-derived riboflavin metabolites to conserved mammalian nucleoside compounds, enabling further investigation of immunosurveillance and immunopathology outside the realm of current clinical paradigms.

## Results

### *M. tuberculosis* and *E.coli* metabolites activate MAIT cells

*E.coli* and *M. tuberculosis* infections stimulate MAIT cells to produce pro-inflammatory cytokines and cytolytic mediators (Le Bourhis et al., 2010; Sharma et al., 2020). From *E.coli*, ribityluracil has been identified as a MAIT cell antigen (Corbett et al., 2014), but the identities and structures of *M. tuberculosis* and mammalian compounds for MAIT cell activation remain unknown. We used *E.coli* as a comparison and *Listeria monocytogenes* (*L. monocytogenes*) as a negative control to determine MAIT cell activation induced by *M. tuberculosis* metabolites through MR1-mediated antigen presentation. Since live *E.coli* (BL21) and *M. tuberculosis* (avirulent H37Ra) strains stimulate MAIT cells in an MR1-dependent manner (Sharma et al., 2020), we sonicated these bacterial cells and obtained small soluble molecules using size exclusion filters with appropriate molecular weight cutoffs (Fig. 1A). Bacterial eluates with overall molecular weight less than 3 kDa were collected and designated as bacterial eluates <3 kDa, because MR1 and ligand binary structures have demonstrated the ligand sizes are typically hundreds of Daltons (Kjer-Nielsen et al., 2012).

**Fig. 1.**
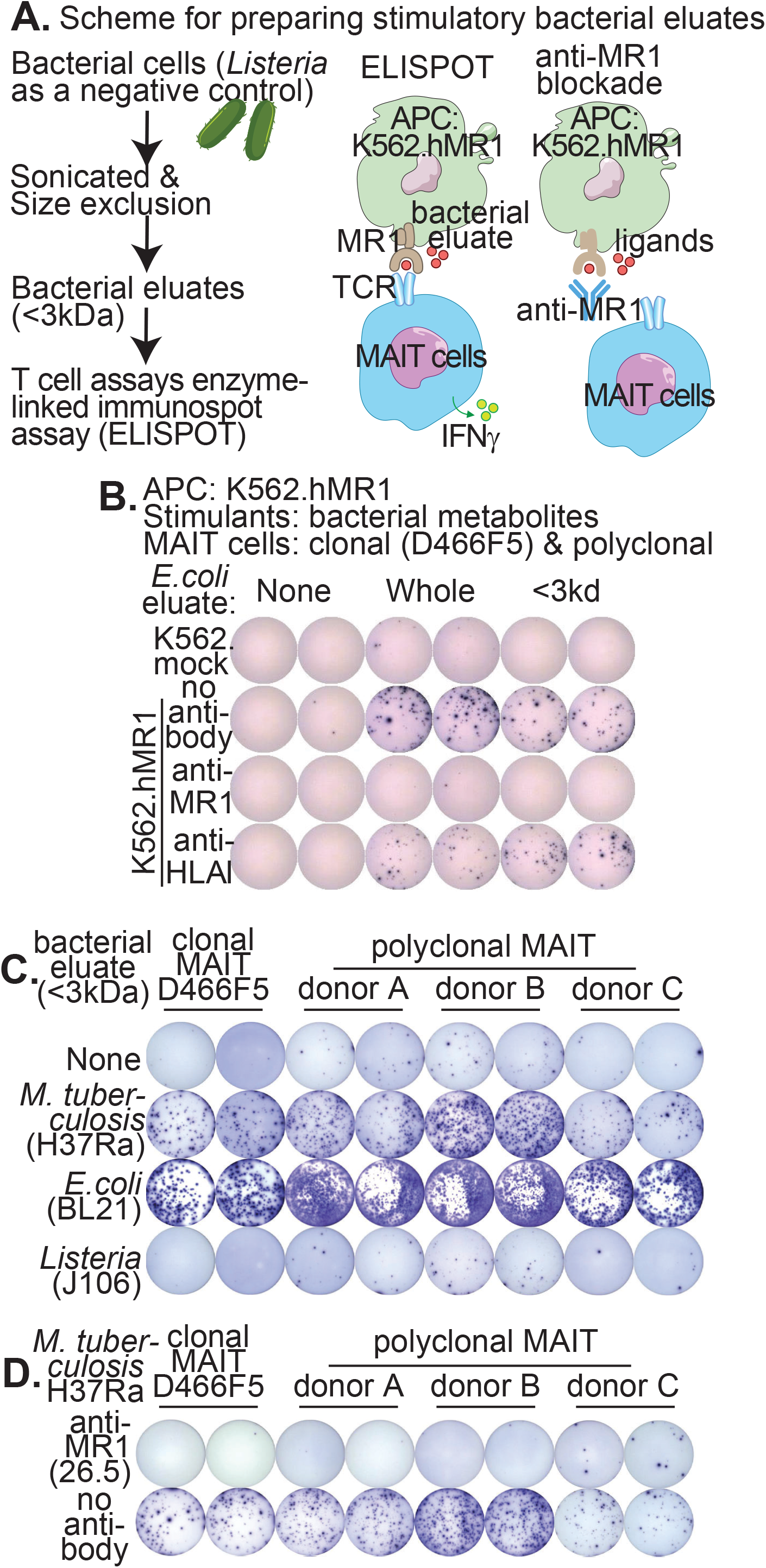
Bacterial eluates activate MAIT cells. Schematic flow shows the procedures to prepare bacterial eluates (<3 kDa) for MAIT cell activation, including the enzyme-linked immunospot (ELISPOT) assay to detect the activated IFNγ^+^ MAIT cell spots. APC: antigen-presenting cells **(A)**. Using *E.coli* eluate (<3 kDa) as an example, crude sonicate and size-filtrated samples are stimulatory for MAIT cells and blocked by the anti-MR1 antibody (26.5, IgG2a). Anti-HLA-I (IgG2a) acts as an isotype control and irrelevant HLA class I protein controls **(B).** *M. tuberculosis* (H37Ra) eluates (<3 kDa) activate the MAIT cells (D466F5) cloned from an active tuberculosis patient, and the polyclonal MAIT cells expanded from the sorted Vα7.2^+^CD161^+^CD4^−^CD8^+^ blood cells of healthy donors. *E.coli* is used as a comparison, and *Listeria* is used as a negative control **(C)**. Clonal and polyclonal MAIT cell activation can be blocked by anti-MR1 antibody **(D)**. Each assay was performed twice independently with similar results.

To test the activity of bacterial eluates (<3 kDa), we incubated human MR1-overexpressing hematopoietic cell line K562 (K562.hMR1) with bacterial eluates (<3 kDa) to stimulate monoclonal and polyclonal human MAIT cells. K562 cells defective in expressing human leukocyte antigens (HLA) minimize the background activity of polyclonal human T cells (de Jong et al., 2010), while K562.hMR1 cells with bacterial infection activate MAIT cells (Sharma et al., 2020). In this study, we used an enzyme-linked immunospot (ELISPOT) approach to measure MAIT cell activation by counting the numbers of interferon γ (IFNγ)-positive spots (IFNγ^+^ spots), which were formed by IFNγ-producing activated MAIT cells (Fig. 1). As a result, both crude sonicated *E.coli* materials and filtered *E.coli* eluates (<3 kDa) activated the MAIT cell line (D466F5), which was cloned from an individual with active tuberculosis (Gold et al., 2010; Sharma et al., 2020). Moreover, MAIT cell activation by bacterial eluate (<3 kDa) was blocked by the anti-MR1 antibody (clone 26.5) (Huang et al., 2005), demonstrating its dependence on MR1-mediated antigen presentation (Fig. 1B). Further, both filtered *E.coli* and *M. tuberculosis* eluates (<3 kDa) activated the MAIT cell line D466F5 and the polyclonal MAIT cells expanded from the blood samples of healthy donors (Fig. 1C). We used *L. monocytogenes* eluate as a negative control (Fig. 1C), because *L. monocytogenes* was previously shown to be unable to activate MAIT cells (Sharma et al., 2020) due to lack of metabolic pathways to produce riboflavin metabolites (Eckle et al., 2015; Kjer-Nielsen et al., 2012). Moreover, polyclonal MAIT cell response to *M. tuberculosis* and *E.coli* appears differently based on donor and strength of response seen in the comparison of donors B and C (Fig. 1C), suggesting differing stimulatory compounds between two bacteria for the stimulation of divergent MAIT cell clonalities. Again, anti-MR1 completely blocked MAIT cell activation by *E.coli* and *M. tuberculosis* eluates (Fig. 1D). Together, the data supported that *M. tuberculosis* activated MAIT cells via MR1 with metabolites that were likely different from *E.coli* metabolites.

### High-pressure liquid chromatography (HPLC) fractionated active *M. tuberculosis* metabolites

To purify *M. tuberculosis* metabolites that activated MAIT cells, we used Shimadzu prominence HPLC to fractionate <3 kDa eluates of *M. tuberculosis* in comparison to *E.coli* and the negative control *L. monocytogenes*. HPLC was performed using a diamond hydride column to run under aqueous normal phase chromatography, which employed a nearly isocratic gradient with a gradually increased polarity. Each injection for HPLC used 75~150 ul bacterial eluates (<3 kDa) with similar total intensities of chromatograms. Bacterial compounds were monitored using ultraviolet (UV) absorbance (Fig. 2A). We collected HPLC fractions of 500ul at a one-minute interval and concentrated them to 50ul using a nitrogen evaporator. The activity of collected HPLC fractions was tested using an ELISPOT assay (Fig. 1A). T cell bioactivity profiles demonstrated that HPLC efficiently separated the stimulatory fractions, which varied between bacteria. For example, we identified *M. tuberculosis* fraction 13 as an active fraction in contrast to the co-eluted inactive *E.coli* fraction 13. Conversely, *E. coli* metabolites showed more rapidly eluting and stronger bioactivity in fractions 6 and 7 compared to *M. tuberculosis*. Overall, data supported the presence of different identities or amounts of compounds in *M. tuberculosis* versus *E. coli* for MAIT cell activation, supporting additional efforts for the structural characterization of potentially novel compounds (Fig. 2B).

**Fig. 2.**
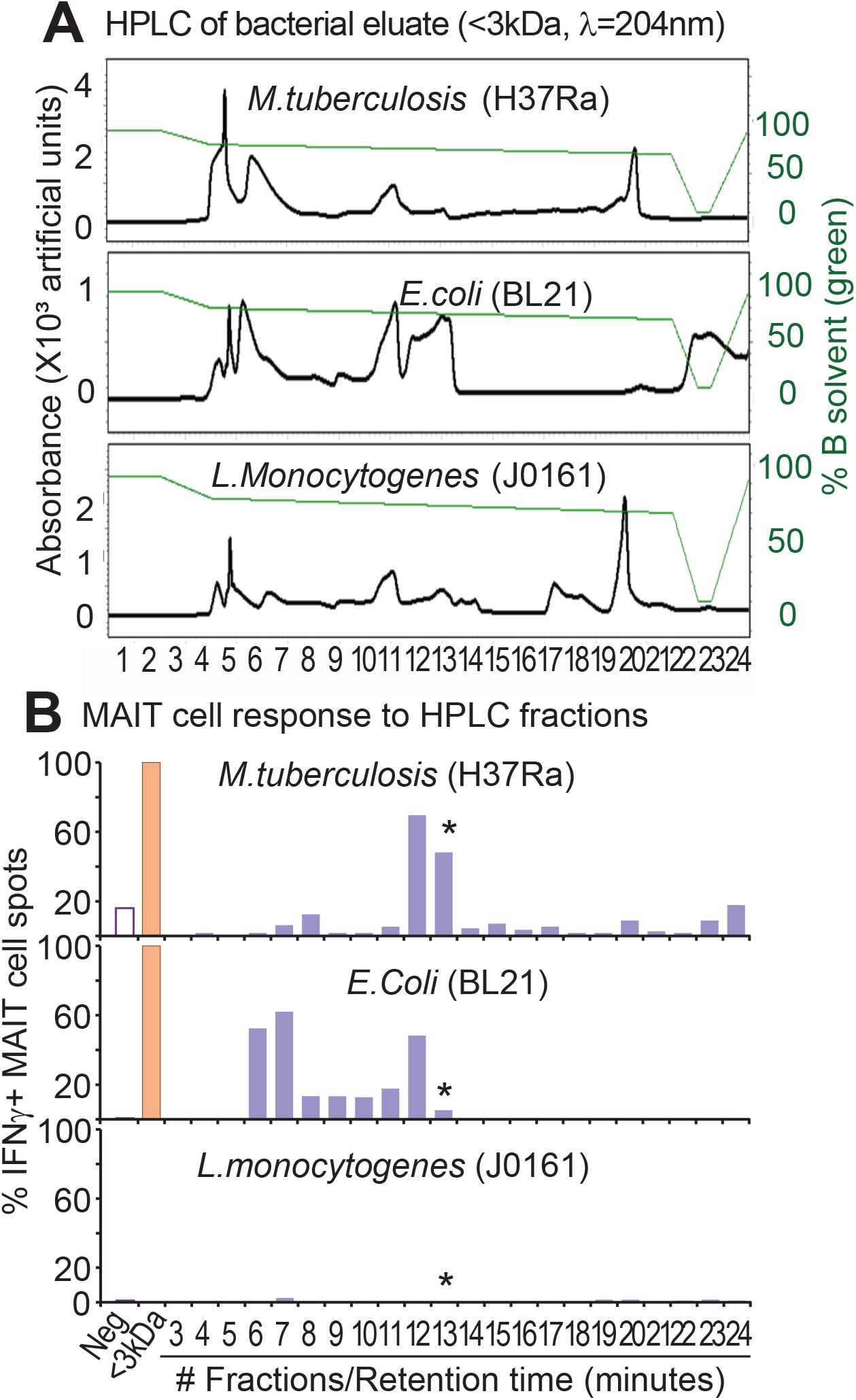
HPLC fractionation of bacterial eluates. HPLC fractionated the bacterial eluates (<3 kDa, 75-150 ul eluate from 1.5-3 ml culture) with a diamond hydride column and an aqueous normal phase gradient (λ, ultraviolet wavelength) **(A).** Collected fractions were concentrated using nitrogen evaporation and confirmed for MAIT cell (D466F5) activation using ELISPOT. Numbers of IFNγ^+^ spots were normalized to positive eluate controls (<3 kDa) from each assay **(B)**.

### Mass spectrometry detected ions enriched in *M. tuberculosis*

Structural prediction of active *M. tuberculosis* metabolites for MAIT cell activation takes advantage of differing patterns in comparative metabolomic profiling of HPLC fractions from multiple bacterial species. Metabolomic profiling of the active versus inactive HPLC fractions concurrently eluted from *M. tuberculosis, E.coli*, and *L. monocytogenes* allow associating the detected ions with bioactivity and rejecting a number of ions present in active *M. tuberculosis* fractions, when the same ions also enriched in non-bioactive fractions from the other two species. In this study, we focused on profiling the active *M. tuberculosis* fraction 13 based on two criteria. First, co-eluted fractions 13 from two control bacteria were nearly inactive and were used as negative controls to show the lists of co-eluted compounds generally irrelevant to bioactivity. Second, fraction 13 was eluted at a mid-range retention time with around 75% solvent B during a gradual enhancement of polarity to avoid artifacts derived from unrelated compounds that accumulated near the solvent front and column regeneration (Fig. 2A).

HPLC-linked mass spectrometry (HPLC-MS) was performed with duplicate runs of each fraction 13 from different bacteria to further separate compounds and generate mass profiles for structural prediction and function confirmation (Fig. 3A). Under the similar aqueous normal phase chromatography using a nearly isocratic gradient, HPLC-MS of fractions 13 resulted in further separation of various compounds and highly overlapping total ion chromatograms for duplicate samples (Figs. 3B and Fig. S1). Mass spectrometry was performed using an Orbitrap Fusion Lumos Tribrid Mass Spectrometer with high sensitivity, and the total ion chromatogram was visualized using the MZmine program (Pluskal et al., 2010). Several major peaks eluted at the early stage and shared by all bacteria, including the negative control *L. monocytogenes*, were considered less relevant to MAIT cell activation. In the middle range of retention, multiple unique base peaks from active *M. tuberculosis* fractions were identified, including a base peak enriched with the ion at a mass-to-charge *(m/z)* value of 291.070 (Fig. 3B and S1). Although results provided rich molecular ions, bioinformatics analyses are needed to comparatively identify enriched or unique ions from *M. tuberculosis* in comparison with *E.coli*. As detailed in Fig. 3A, we performed clustering and peak picking, respectively, to comprehensively identify molecular ions and features enriched in the active fraction. Chromatic and spectral visualization confirmed individual molecular ions enriched in the active fraction. Fragmentation of targeted molecular ions predicted candidate compounds for functional confirmation (Fig. 3A).

**Fig. 3.**
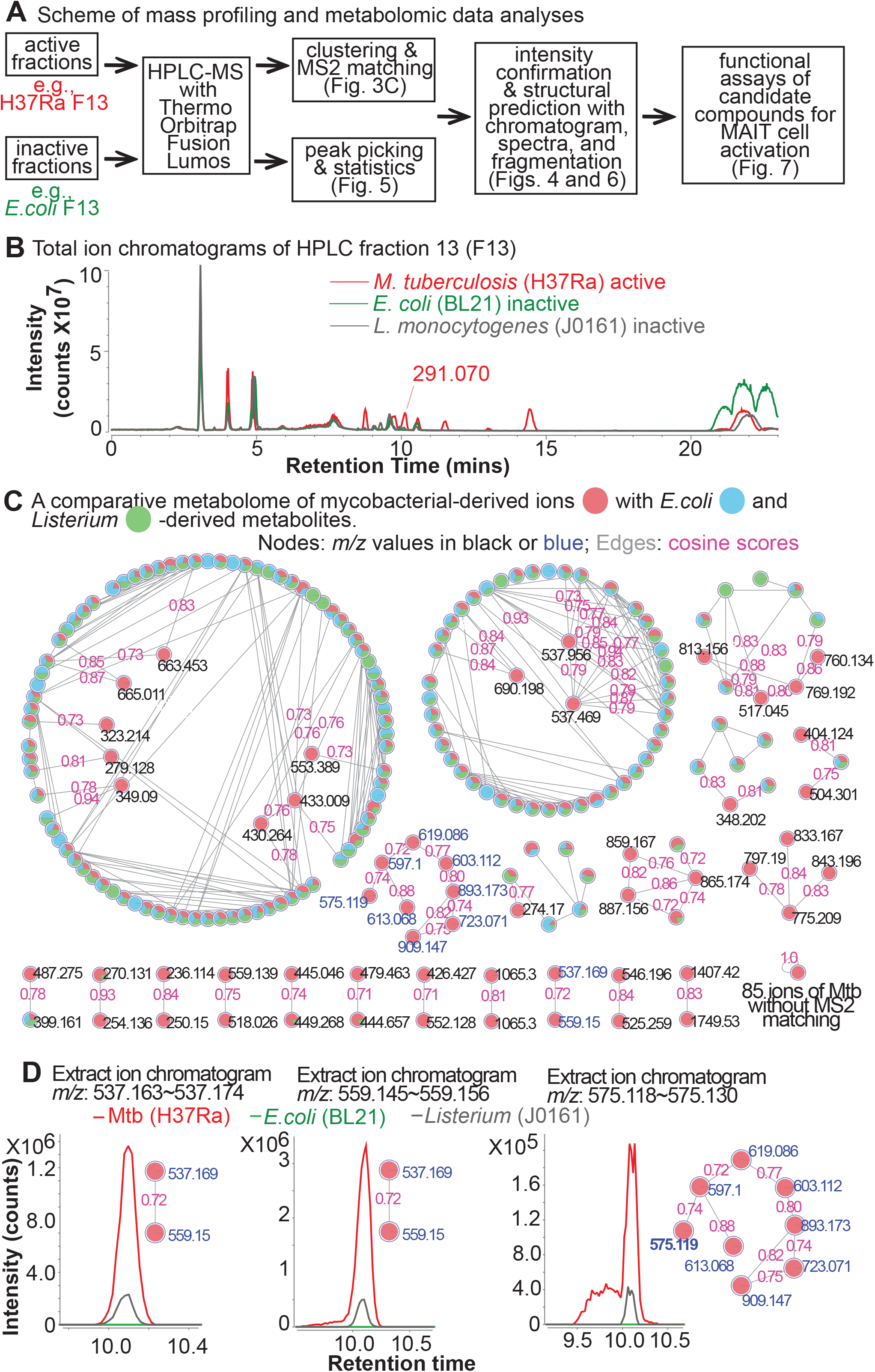
Comparative metabolomics and clustering of mass profiles. Schematic flow of functional comparative metabolomics combines chemical, computational, and functional analyses **(A)**. The total ion chromatogram from duplicated runs of each bacterium is overlaid to show unique base peaks from *M. tuberculosis* **(B)**. GNPS clusters the molecular ions differentially shared and enriched among three bacteria. *M. tuberculosis*-associated ions are displayed with colors annotating the relative association of molecular ions with each bacterium. Edges with cosine scores represent the similarity of fragmentation patterns for ions **(C)**. Extract ion chromatograms of multiple clustered molecular features (highlighted in blue) enriched in *M. tuberculosis* **(D)**.

### GNPS clustered multiple molecular ions with inosine moiety

Global Natural Product Social Molecular Networking (GNPS) provides libraries of known compounds and algorithms of spectral matching that facilitate the identification of natural products based on tandem mass spectrometry profiles (Olivon et al., 2017). We input mass profiles to the GNPS program and performed a molecular network search based on the default setting (Wang et al., 2016) to assign molecular ions enriched with *M. tuberculosis* and estimate the similarity of fragmentation pattern using cosine scores (Fig. 3C). As a result, GNPS accurately assigned *m/z* values, suggested the molecular ions enriched in different samples, and provided a network based on similar fragmentation patterns measured by cosine scores. A significant match was measured by cosine scores higher than 0.7 (Nothias et al., 2020). However, it did not yield a comprehensive prediction of various molecular identities by matching our fragmentation profiles with the available fragmentation profiles in the database using a molecular library search module.

By examining ion clusters more enriched in *M. tuberculosis* (Fig. 3C), we manually extracted ion chromatograms of multiple molecular ions from two clusters to determine whether their higher intensity from *M. tuberculosis* samples was valid. We further examined the fragmentation patterns of these molecular ions as a result of collision-induced dissociation-mass spectrometry (CID-MS) and identified a neutral loss of the mass 268.081 from multiple molecular ions (Fig. S2), which shared high cosine scores (>0.7) implicating significantly matched fragmentation (Fig. 3D). Since multiple collisional spectra showed the neutral loss and ion forms of the mass 268.081 (Fig. 3D), we sought to predict its protonated and sodiated ions at *m/z* of 269.088 and 291.070, respectively. The signature fragment of protonated hypoxanthine *(m/z* 137.046 Fig. 4AB) and sodiated hypoxanthine (*m/z* 159.028 not shown) supported the prediction of inosine, ribosylhypoxanthine, or isomers thereof, in comparison to the collision pattern of a synthetic inosine as a reference (Fig. 4CD). Although this purine species enriched in *M. tuberculosis* (Fig. 4), it was also eluted from *L. monocytogenes* at a lower intensity and considered a conserved molecule in various bacterial and mammalian species (Fig. 4A). Therefore, inosine and isomers are unlikely to explain the MAIT cell stimulation from the *M. tuberculosis* eluate and HPLC fraction seen in our bioassays (Figs. 1 and 2). However, the prediction of inosine or isomers suggests that nucleoside moieties are likely enriched in this *M. tuberculosis* sample and potentially contain MAIT cell regulators to be identified using statistical annotation and functional tests. Using ELISPOT assays (Fig. 4E), we confirmed that inosine or arabinosylhypoxanthine could weakly stimulate an MAIT cell clone (Fig. 4F) and polyclonal MAIT cells (Fig. 4G). Although some nucleosides, such as inosine, were more detected in *M. tuberculosis* than in *E.coli* (Rizvi et al., 2019), inosine is not *M. tuberculosis-specific* and a weak MAIT cell stimulation by inosine could not interpret the activity of *M. tuberculosis* fraction 13. Nonetheless, these data still strongly suggests nucleoside compounds with various structural modifications likely bind to MR1 and regulate MAIT cell responses.

**Fig. 4.**
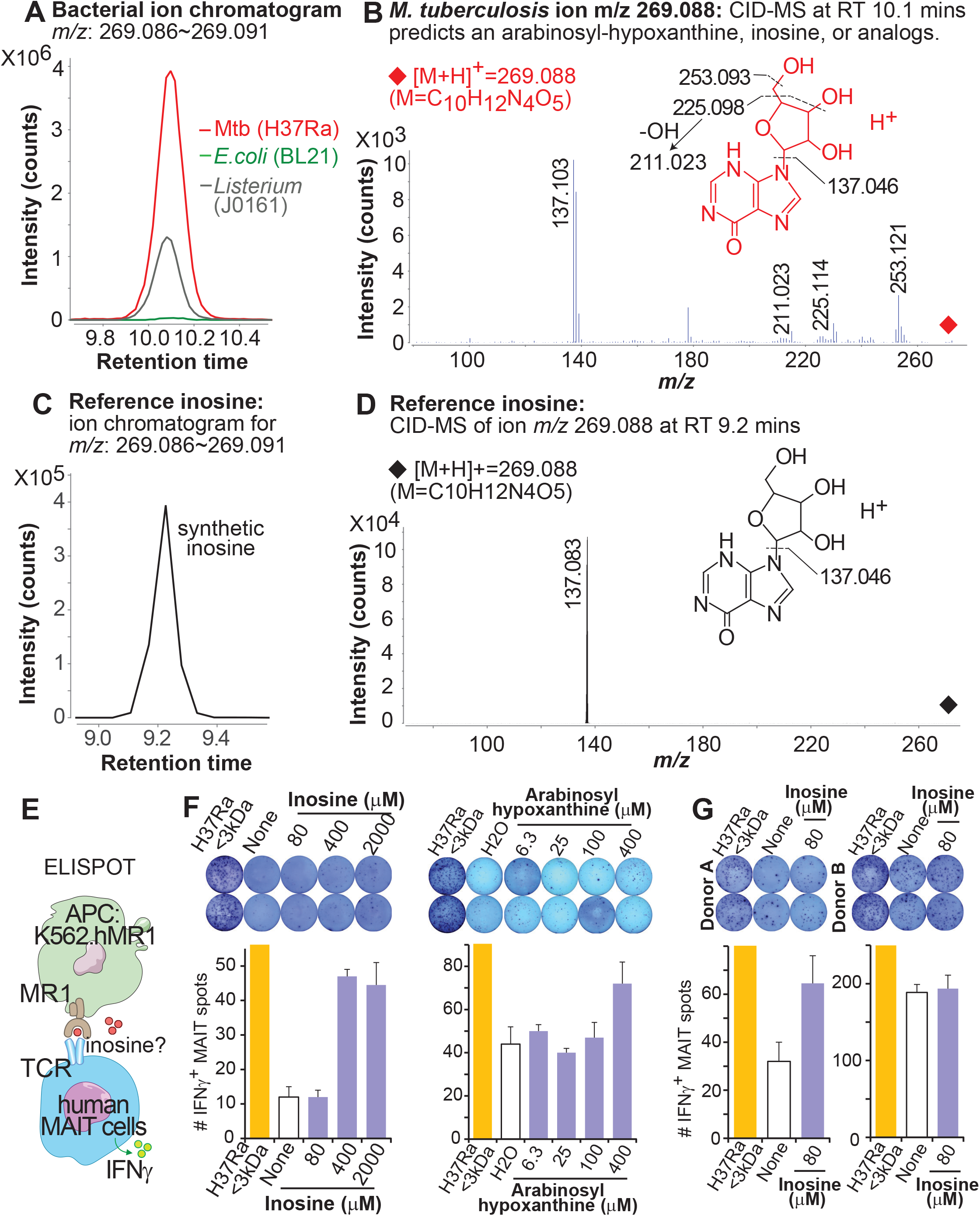
Prediction and functional tests of inosine or analogs enriched in M. tuberculosis. The ion chromatogram **(A)** and collisional spectrum **(B)** of the *M. tuberculosis-enriched* parental ion *m/z* 269.088 were detected using LC-MS and collision-induced dissociation-mass spectrometry (CID-MS), in comparison to the ion chromatogram **(C)** and collisional spectrum **(D)** of reference compound inosine. An ELISPOT assay **(E)** was used to test the role of inosine and isomer in the activation of clonal MAIT cells **(F).** Polyclonal MAIT cells expanded from the sorted Vα7.2^+^CD161^+^CD4^−^ CD8^+^ blood cells of healthy donors were also tested with response to inosine **(G).** The ELISPOT data are from one of two independent assays with similar results.

### XCMS program clustered the molecular features enriched in *M. tuberculosis*

To statistically and more comprehensively quantify the enriched molecular features from *M. tuberculosis* fractions, we used the XCMS program (Tautenhahn et al., 2012) to de-convolute mass spectra from duplicate runs of HPLC fraction 13 of *M. tuberculosis, E.coli*, and *L. monocytogenes*, together with the solvent blank. XCMS running on an R platform can pick peaks, calculate intensity counts, and align retention times to define molecular features (Huang et al., 2011; Layre et al., 2011) by generating accurate mass, retention times, and intensity values for each detected ion (Fig. 5A). From these results, we defined the molecular features enriched with *M. tuberculosis* using the following inclusion criteria: ***(i)*** relatively low background intensities in the solvent blank (<10^6^ counts), ***(ii)*** above minimal intensities from *M. tuberculosis* (>10^5^ counts), ***(iii)*** higher observed intensities in *M. tuberculosis* than in solvent blank (>50-fold) and in *L. monocytogenes* or *E. coli* (>5-fold) (Fig. 5B). Upon analysis and inclusion with XCMS and above criteria, we further manually examined the chromatogram and spectra of the obtained molecular features (n=823). Results confirmed that XCMS program assigned accurate *m/z* values (generally <10 parts per million, ppm), which are critical for formula and structure prediction. However, XCMS program picked both molecular and isotopic ions and provided sensitive quantification of weakly detected ions. To obtain molecular ions differentially expressed in *M. tuberculosis* and *E. coli*, we removed a high percentage of molecular features representing isotopic ions (24.4%), with weak chromatogram, mass spectra, or missing isotopic patterns (44.5%), or without visual difference of chromatogram (12.5%). We then obtained 170 molecular features to show the statistical tests using analysis of variance (ANOVA) for cross-bacterial differential expression and a volcano plot displayed the molecular features with 5-fold or 100-fold higher intensity in *M. tuberculosis* versus *E.coli* with a significant p-value (<0.05) (Fig. 5B). Results defined more than a hundred of molecular features (n=146) enriched in *M. tuberculosis* instead of *E. coli* (Fig. 5B). Therefore, the comparative metabolomics with peak picking, accurate mass assignment, intensity quantification, statistical tests, and volcano plot visualization provides the molecular features differentially associated with bacteria and bioactivity for further structural and functional characterization.

**Fig. 5.**
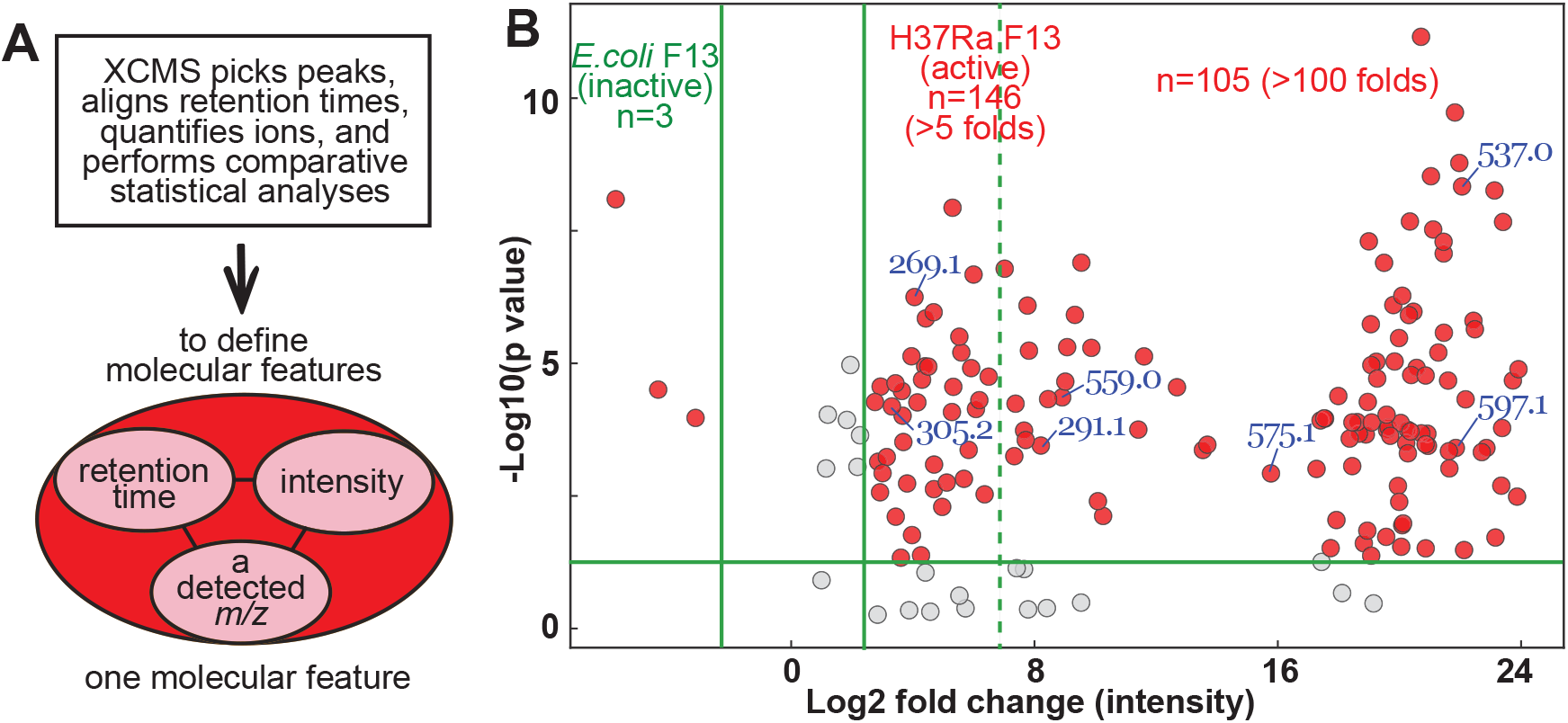
Statistical analyses of molecular features enriched in M. tuberculosis. Mass profiles of duplicated runs for each bacterium were analyzed using the XCMS program to generate molecular features. A molecular feature is used to define a mass spectral peak from precursor ion spectral profiles with retention time, intensity, and a detected mass-to-charge (*m/z*) value **(A)**. A volcano plot shows the molecular features enriched in *M. tuberculosis* upon confirmation with ion chromatograms and mass spectra. Vertical solid green lines show a 5-fold cutoff of intensity; a vertical dash green line shows a 100-fold cutoff; and the horizontal green line shows a p-value of 0.05 **(B).**

### Prediction of methylinosine and formyluridine analogs enriched in *M. tuberculosis*

Upon further annotation of molecular features enriched with *M. tuberculosis* in the volcano plot (Fig. 5B), many molecular features with relatively small *m/z* values, including 269.1, 291.1, 305.2, and 353.0, fall in the range of mass values of reported MR1 ligands (Corbett et al., 2014; Harriff et al., 2018; Kjer-Nielsen et al., 2012), and are generally above 5-fold cutoff. Thus, the volcano plot displays the molecular features enriched in *M. tuberculosis* as candidate targets for further structural prediction. Technically, mass spectrometry accurately detected parental molecular ions within around 5 ppm, as shown by multiple tested results of the same parental ions in a mass spectrometry run. Mass values of fragments from CID-MS profiling were variable with <0.1 mass unit difference among multiple collisions of the same parental molecular ions. But these results still provided valuable predictions of candidate ions. Beyond the prediction of inosine and analogs (Fig. 4), we found multiple molecular ions matching various species of modified nucleosides. The molecular ion *m/z* 305.086 enriches with *M. tuberculosis* (Fig. 6A) and matches the mass of sodiated methyl-inosine *(m/z* 305.086) at very high accuracy (<1 ppm). Collisional spectra with multiple weak fragments match the collision of methylinosine, including a typical sodiated hypoxanthine fragment at *m/z* 143.029 (Fig. 6B), which was predicted as the sodiated hypoxanthine losing a hydroxyl group and is different from a synthetic 1-methylinosine with a methylated hypoxanthine at *m/z* 173.043 (Fig. 6D). Thus, the precursor ion *m/z* 305.086 can be predicted as a methylinosine analog with a ribosyl methylation, which commonly locates at the 2’-OH position of ribose (Ayadi et al., 2019; Drazkowska et al., 2022). The synthetic reference 1-methylinosine compound with methylation at the N1 position of the purine base was detected at a similar retention time to the predicted methylinosine analog (Fig. 6CD) and displayed a fragment of a sodiated methyl-hypoxanthine (Fig. 6D), supporting a structural similarity to and yet difference from the predicted methylinosine analog. Collision of synthetic guanosine also released the protonated guanine at *m/z* 152.057 (Fig. 6EF), providing an alternative control for methylinosine. Furthermore, a molecular ion *m/z* 353.038 enriched with *M. tuberculosis* (Fig. 6GH) is predicted as formyluridine monophosphate with fragments indicating the losses of a hydroxyl group *(m/z* 335.027) or two carbonyl oxygens *(m/z* 307.030) (Fig. 6H). The reference synthetic compound deoxyformyluridine showed an early eluted ion chromatogram (Fig. 6I) due to more hydrophobic nature without a phosphate group and a similar but a more completed collisional spectra in a separate run using targeted analysis (Fig. 6J). In summary, our mass spectrometry analyses predicted methylated and formylated nucleoside compounds, such as methylinosine and formyluridine, enriched in the *M. tuberculosis* fraction.

**Fig. 6.**
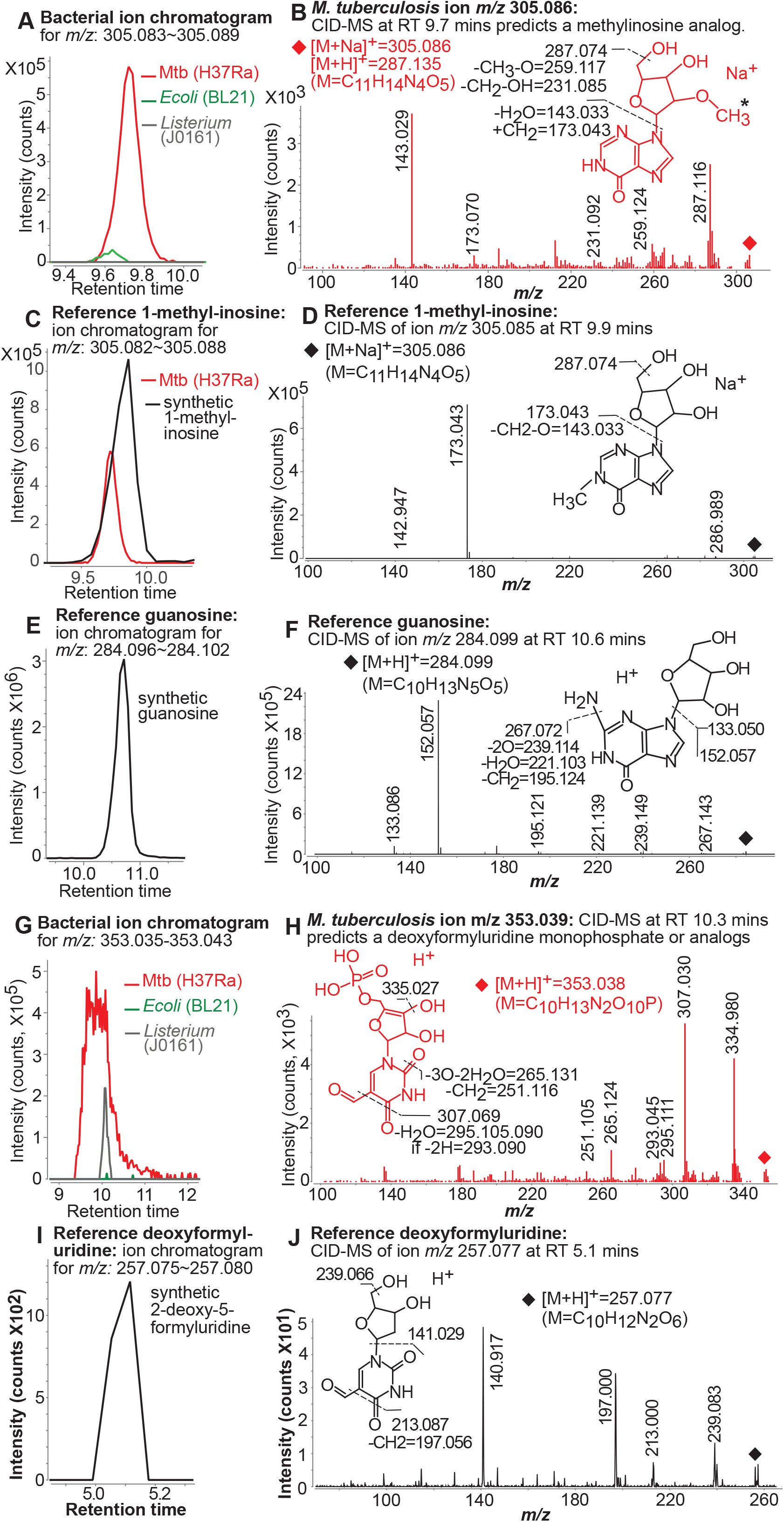
Prediction of nucleosides or analogs enriched in M. tuberculosis. The ion chromatogram **(A)** and collisional spectrum **(B)** of the *M. tuberculosis-enriched* precursor ion of *m/z* 305.086 were compared with the ion chromatogram **(C)** and collisional spectrum **(D)** of the synthetic compound 1-methylinosine, and the ion chromatogram **(E)** and collisional spectrum **(F)** of the synthetic compound guanosine. Further, the ion chromatogram **(G)** and collisional spectrum **(H)** of the *M. tuberculosis-enriched* precursor ion of *m/z* 353.039 were compared with the ion chromatogram **(I)** and collisional spectrum **(J)** of the synthetic compound deoxyformyluridine.

### Autodock analyses suggested the potential of multiple nucleosides for MR1 binding

Although the predictions of nucleoside compounds enriched in *M. tuberculosis* fraction were mainly based on high accuracy matches of parental ions but low accuracy matches in fragmentation, results remain valuable to suggest nucleoside analogs for MAIT cell functional tests. To select which nucleoside metabolites were likely more promising in further functional assays, recently developed computational program Autodock Vina with deep machine learning modules for metabolite docking can be used to assess the binding compacity of candidate ligands with MR1 protein (Eberhardt et al., 2021; Liu et al., 2020). This supplemental step is highly meaningful because multiple predicted *M. tuberculosis* nucleosides are unavailable in synthetic forms and the available nucleoside analogs are not *M. tuberculosis*-specific but conserved in mammals. We selected four MR1 crystal structures with or without covalent bounding with stimulatory or inhibitory ligands. Binding compacity was measured by the absolute values of free energy (Vina scores, Fig. S3). A threshold was set based on the comparison of MR1 docking with the known MR1 ligands or analogs, and the non-ligand metabolites of Vitamin B family. As a result, multiple nucleoside metabolites, such as guanosine and 2-O-methylinosine, achieved an absolute Vina score higher than the threshold, suggesting a high predicted affinity to bind to MR1 protein and a potential for regulating MAIT cell responses.

### Guanosine activated and deoxyformyluridine inhibited MAIT cell responses

Whereas our data predict some nucleosides, such as 2-O-methylinosine (Fig. 6AB) and dimethyl-amino-guanosine (not shown), are more abundant in *M. tuberculosis*, these compounds were not available. However, the synthetic analogous 1-methylinosine and N2-dimethylguanosine were unable to activate MAIT cells in multiple assays (data not shown), suggesting that the dual methylation at the N2 amino position potentially depleted the activity. Interestingly, we could demonstrate the MAIT cell-regulation activity of their nucleoside analogs, including guanosine and deoxyformyluridine as conserved compounds in microbes and mammals. Guanosine activated clonal (Fig. 7A) and polyclonal (Fig. 7B) human MAIT cells at relatively high concentrations, such as 400 uM to 2 mM. MAIT cell responses stimulated by guanosine can be blocked by MR1 ligand acetyl-6-formylpterin (Ac-6-FP) as MAIT cell antagonist and by anti-MR1 antibody, supporting its dependence on MR1-mediated antigen presentation (Fig. 7C). Surprisingly, 2-deoxy-5-formyluridine, simply called deoxyformyluridine in this study, clearly inhibited MAIT cell activation that were induced by *M. tuberculosis* eluate (<3 kDa) in both clonal (Fig. 7D) and polyclonal (Fig. 7E) human MAIT cells. The deoxyformyluridine is a nucleoside compound conserved in both bacteria and mammalian cells (Kim and Almo, 2013; McCown et al., 2020). The inhibitory role of deoxyformyluridine likely provides a negative feedback control in MAIT cell responses and a conserved regulatory role of MAIT cells in mammalian cells. Together, we identified the nucleoside guanosine that activated MAIT cells dependent on MR1-mediated antigen presentation and the nucleoside deoxyformyluridine that inhibited MAIT cell responses. We speculated that these conserved nucleoside metabolites potentially regulate MAIT cell responses in infected or uninfected conditions.

**Fig. 7.**
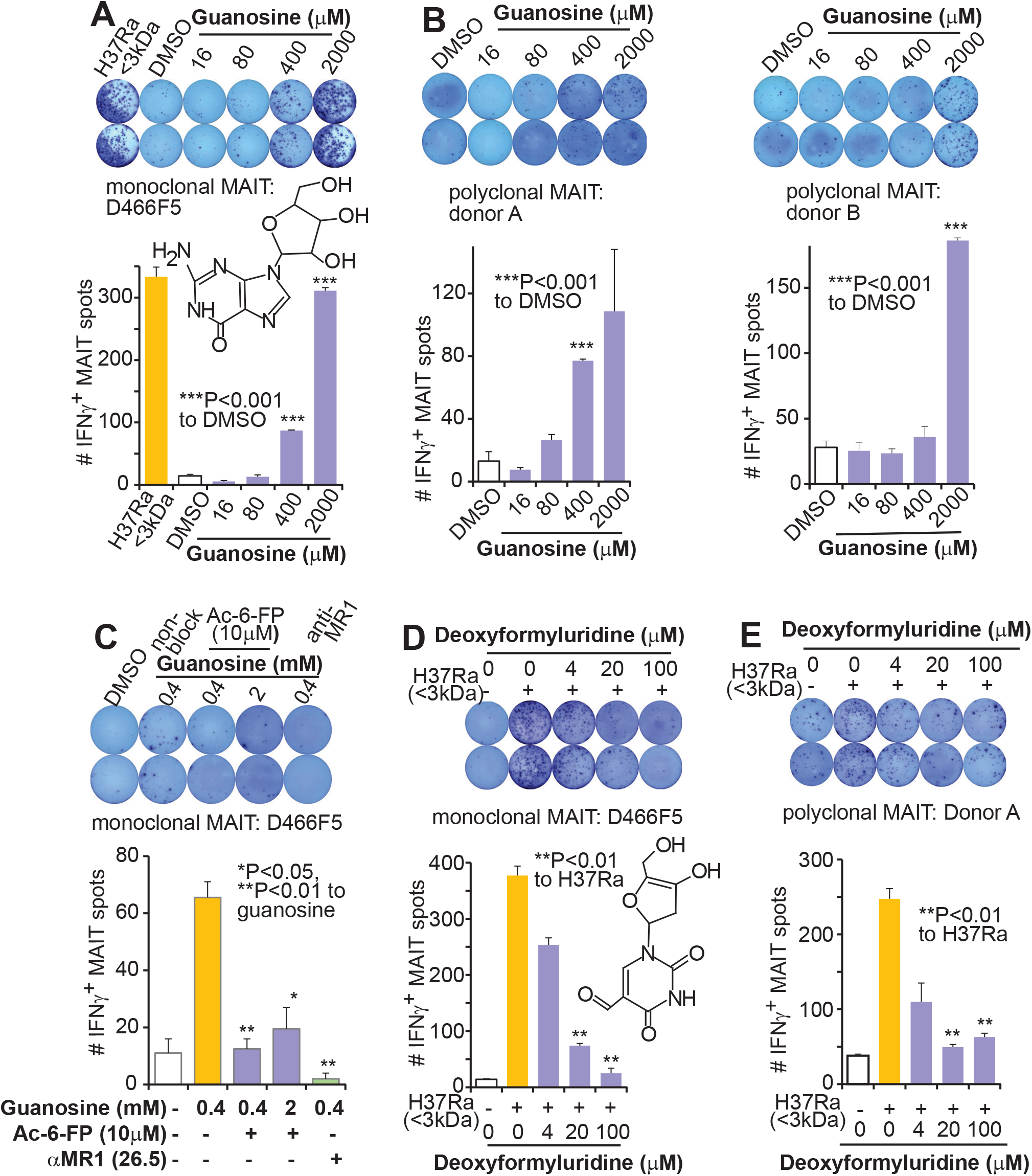
MAIT cell activation and inhibition with nucleoside compounds. ELISPOT measures the clonal **(A)** and polyclonal **(B)** MAIT cell responses to the stimulation of guanosine. The polyclonal MAIT cells were expanded from the sorted Vα7.2^+^CD161^+^CD4^−^CD8^+^ blood cells of healthy donors. MAIT cell stimulation by guanosine is blocked by MAIT cell antagonist acetyl-6-formylpterin (Ac-6-FP) and anti-MR1 antibody (clone 26.5) **(C).** Deoxyformyluridine differentially blocks H37Ra to stimulate the clonal **(D)** and polyclonal **(E)** MAIT cell responses. P-values were calculated using an unpaired t-test in comparison to negative controls (blank columns) in stimulating assays and to positive controls (yellow columns) in blocking assays. Significance considers one to three comparison tests with p<0.05, 0.025, and 0.01, respectively. Data are from one of at least two independent assays with similar results.

## Discussion

The functional metabolomics in this study combined functional monitoring with consecutive steps of chemical purification and metabolomic profiling to build a comparative platform for identifying metabolites and natural products with specific biological roles. We monitored the bioactivity of unknown compounds throughout the purification process, starting from size-based filtration to achieve bioactive bacterial eluates (<3 kDa), polarity-based HPLC separations of active eluates with the inactive as a control, and further activity monitoring of HPLC fractions, structural prediction of molecular ions from bioactive HPLC fractions, to eventual functional tests of candidate compounds. This consecutive functional tracking of the targeted samples and candidate compounds allowed us to determine whether the annotated molecular features and predicted compounds contribute to bioactivity. Comparative metabolomic profile analyses supported by bioinformatics tools is an essential step in this platform to separate and predict the candidate molecular features enriched or unique in bioactive *M. tuberculosis* samples. Further, extract ion chromatogram and mass spectra can accurately confirm the presence of targeted molecular ions with bioactive samples, providing candidate targets for structural and functional characterization. Moreover, automated or targeted collisional analyses generated dissociated fragments that either predicted the identities of a full molecule or structural moieties as a part of a molecule, such as neutral loss of an inosine moiety or fragments of purine or pyrimidine moieties, facilitating the selection of candidate compounds or alternative analogs for functional assays. Using Autodock Vina analysis with incorporated deep machine learning (Eberhardt et al., 2021; Liu et al., 2020), we further predict potential binding capacity of candidate metabolites with MR1 protein. We also selected potential candidates for MAIT cell activation tests by further considering their overall reported molecular sizes and structural moieties critically for MR1 binding and TCR interaction (Corbett et al., 2014; Kjer-Nielsen et al., 2012; Patel et al., 2013; Rossjohn et al., 2015). Together, this functional metabolomics integrates functional monitoring of differentially purified natural products and comparative annotation of molecular features associated with activity, leading to structural and functional elucidation of small metabolites and natural products from complex biological samples for immune regulation.

Previously reported bacterial antigens for MAIT cells show a common feature in chemical structures, which usually comprise ring structures and ribityl moieties. These bacterial antigens, ribityllumazines, ribityluracils, and photo-lumazine, are intermediates in microbial riboflavin metabolism and were identified from *Salmonella typhimurium, E.coli*, and *Mycobacterium smegmatis* (Corbett et al., 2014))(Harriff et al., 2018; Kjer-Nielsen et al., 2012), respectively. These bacterial antigens stimulate MAIT cell responses strongly and are considered “potent” MAIT cell antigens. Different from intermediate metabolites in riboflavin metabolism, the nucleosides guanosine and deoxyformyluridine reported in this study represent the compounds shared between bacteria and mammalian cells, and therefore represent new candidates for positive or negative regulation of MAIT response in either infected or uninfected hosts. Indeed, results from multiple recent studies support the existence of uncharacterized MAIT cell antigens likely different from riboflavin metabolites. Divergent MAIT cell clones respond differently to Gram-positive versus Gram-negative bacteria, including species of *Mycobacterium*, *Salmonella*, and *Staphylococcus* (Meermeier et al., 2016), supporting pathogen selectivity and ligand discrimination of MAIT cell clones. Similarly, MAIT cells with diverse TCRβ chains and limited variation of the fragment J of TCRα chain respond differently to the same bacteria and the same MAIT cell clones respond differently to various bacteria, suggesting the expression of different identities or quantities of antigens (Gold et al., 2014; Harriff et al., 2018). Responses to uncharacterized antigens different from riboflavin metabolites are also associated with non-canonical TCR expression of MAIT cells (Crowther et al., 2020; Lepore et al., 2017; Meermeier et al., 2016). Moreover, HPLC retained active *M. tuberculosis* fractions different from *E.coli* fractions studied here. This evidence together supports antigen diversity across various bacteria and existence of novel compounds for MAIT cell activation. The finding of nucleoside compounds in this study unlikely explains the above heterogeneous MAIT cell responses, but provides natural products different from previously reported riboflavin metabolites to modulate MAIT cell reactivity.

Although unique *M. tuberculosis* metabolites for MAIT cell activation remain to be elucidated, we have identified in this study the bioactive nucleoside compounds that are conserved in bacterial and mammalian cells, unlike bacterial-derived riboflavin metabolites. Endogenous mammalian ligands have been implicated in multiple sterile or uninfected conditions, such as the *in vitro* autoreactivity of MAIT cells from mice (Huang et al., 2005; Huang et al., 2008; Le Bourhis et al., 2010), blockade of mouse MAIT cell responses using an antagonist MR1 ligand (Toubal et al., 2020), and human MAIT cell stimulation without exogenous compounds (Lett et al., 2022). In human inflammatory conditions, unidentified circulating metabolites of unknown bacterial or mammalian sources can activate the MAIT cells isolated from the inflammatory sites, such as the liver (Lett et al., 2022). Either these intracellular stimulants from antigen-presenting cells or extracellular circulating stimuli in human blood remain elusive (Lett et al., 2022). A weak stimulator such as guanosine and an inhibitor such as deoxyformyluridine nucleosides tested in this study can potentially function as conserved candidate mammalian or bacterial compounds of intracellular and extracellular sources for regulating MAIT cell response.

Nucleosides are generally conserved compounds in viruses, bacteria, and animals for nucleic acid synthesis and as potential targets for drug design (Diab et al., 2007; Mashabela et al., 2019). Regarding the effective concentration of exogenously supplied guanosine for MAIT cell activation, it remains unknown how efficient the extracellular guanosine can be delivered to intracellular compartments for MR1 ligand loading since endocytic antigen loading is crucial for MAIT cell activation (Harriff et al., 2016; Huang et al., 2008). Nonetheless, nucleosides are hydrophilic, thus difficult to cross hydrophobic plasma membranes (Diab et al., 2007). Considering that liposomes have successfully facilitated the intracellular and endocytic delivery of nucleoside compounds as in nucleoside analog delivery for cancer therapy (Diab et al., 2007) and RNA vaccine delivery to fight viral infections (Hou et al., 2021), we improved MAIT cell activation by guanosine using commercial liposomes to potentially facilitate the intracellular delivery of guanosine (Fig. 7). Therefore, the high concentration of guanosine needed for MAIT cell activation is partially due to unknown efficient approach to deliver extracellular guanosine to intracellular compartments. However, the exact intracellular or endocytic concentration of guanosine for MR1 loading requires further accurate measurement in future studies. If considering the weak autoreactivity of human MAIT cells in physiological conditions, it is likely that basal intracellular concentration of guanosine in mammalian cells such as human macrophage remains insufficient to stimulate MAIT cells. Nucleoside metabolic pathways, by which free bases, nucleosides, and nucleotides are interconverted and contribute to highly conserved metabolic processes of nucleic acids and proteins, thus, are considered critical to maintaining the intracellular level of guanosine. These intervening metabolic pathways will certainly complicate further targeted quantitative measurement of intracellular guanosine concentrations in health and disease conditions.

Functionally, guanosine has been reported to mediate other cellular responses of T cells unlikely involving MR1 molecule or MAIT cell activation. For example, guanosine reduces the production of inflammatory cytokines and inhibits NF-κB signaling pathways as an anti-inflammatory agent (Jackson and Mi, 2014; Luo et al., 2021). Similarly, another purine nucleoside, adenosine, has been suggested to directly inhibit T cell responses by interacting with adenosine receptors on T cells. In particular, adenosine suppresses CD8^+^ T cell responses by inhibiting TCR signaling (Linnemann et al., 2009), and blockade of the adenosine receptor in T cells enhances CD8^+^ T cell responses (Beavis et al., 2017). We have considered the exclusion of these off-target effects of nucleosides in MAIT cell activation. To confirm the MR1-dependency of guanosine for MAIT cell activation and exclude the possibility of guanosine interaction with adenosine or other receptors in MAIT cells, we applied an anti-MR1 antibody and antagonist ligand (Ac-6-FP) to block the predicted MR1 ligand binding with guanosine, resulting the blockade of MAIT cell activation and supporting MR1-dependence. Regarding methylated nucleoside compounds, we tested dimethylguanosine and 1-methylinosine that were unable to stimulate MAIT cells. Single methylation at the exocyclic amino group of guanosine can be a common modification in RNA and conserved in bacterial and eukaryotic RNA (Pallan et al., 2008). The dual methylation likely eliminates the ability of the amino group function to donate electrons in hydrogen bonds and alters its pairing behavior in transfer RNA or ribosomal RNA formation. However, dimethylamino-guanosine and 2-O-methylinosine as predicted from the *M. tuberculosis* fraction are unavailable for functional assays. These modified forms of nucleosides remain conserved and important in biological processes. Methylation of inosine is also conserved, for example, methylinosine analogs can be detected in mycobacteria (Kavalchuk et al., 2022) and human urine (Roux et al., 2012). More interestingly, the inhibitory compound deoxyformyluridine is a formyl-containing nucleoside that is created under oxidative stress in differentiated cells (Runtsch et al., 2021). Our finding suggests that an inhibited MAIT cell response can likely be induced under the oxidative stress condition of MR1-expressing cells as a feedback regulation since cytokine production upon MAIT cell activation is expected to induce oxidative stress in phagocytes (Ellis and Beaman, 2004). In summary, our findings of stimulatory activity of guanosine dependent on MR1-mediated antigen presentation and inhibitory activity of deoxyformyluridine in MAIT cell responses reveal that conserved nucleoside compounds from bacterial and mammalian cells can stimulate or inhibit MAIT cell responses, potentially regulate MAIT cell functions in infectious and inflammatory conditions.

## Materials and Methods

### Generation of bacterial eluates

*Mycobacterium tuberculosis (M. tuberculosis*, avirulent strain H37Ra, Colorado State University, Fort Collins, CO) (Roura-Mir et al., 2005) was cultured for 5 to 6 days in middlebrook 7H9 complete medium at 37 °C using an orbital shaker with a speed setting at 270 rpm. *Escherichia coli (E. coli* strain BL21, New England BioLabs) and *Listeria monocytogenes (L. monocytogenes* strain J0161, Bei resources) were cultured overnight at 37 °C in the Luria-Bertani broth using an orbital shaker at 100 rpm. Bacteria were harvested at a late log growth phase, washed with phosphate buffer saline (PBS), and measured for their absorbance (optical density at wavelength 600 nanometres, OD_600_) according to the report (Biesta-Peters et al., 2010). OD_600_ provides a semi-quantitative method to estimate bacterial cell numbers that were tested and sufficient for MAIT cell activation. Bacterial culture was harvested while the result of OD_600_ was around 1, washed twice with 0.9% sodium chloride, and re-suspended the bacterial pellet from 1.6 liter culture in 40ml (40:1) Optima H2O (Fisher Brand). The bacterial suspension was sonicated using a sonicator with an intensity setting of 50% for 1-min sonication and 1-min interval. For thorough cell disruption, sonication was performed 30 times in an ice bath for *M. tuberculosis* and 10 times for *E.coli* or *L. monocytogenes*. Sonicated mixture was run through serial centrifugation units with Ultracel regenerated cellulose membrane in a range of molecular weight cutoffs (Vivaspin or GE Healthcare) from 100k to 3k Daltons. The bacterial metabolite eluates from the final 3k Dalton cutoff were collected (designated bacterial eluate <3 kDa) and tested for MAIT cell activation.

### Generation of human MR1-expressing cells and human MAIT cells

We overexpressed the human MR1 protein by transducing the wild-type human MR1 gene using retroviral vector pMXIP (Huang et al., 2009) into human HLA-defective myeloid cell line K562 (de Jong et al., 2010; Roder et al., 1979) as we reported (Huang et al., 2005; Sharma et al., 2020). The transduced K562.hMR1 cells were selected under 5 ug/ml puromycin or sorted using flow cytometry upon anti-MR1 staining (clone 26.5) for high expression of the recombinant human MR1 protein. K562.hMR1 cells were then expanded and saved as aliquots at the cryogenic storage condition. Monoclonal MAIT cells (D466F5) from an active tuberculosis patient were reported previously (Gold et al., 2010) and expanded in our lab using a routine T cell expansion protocol as we reported (Huang et al., 2011; Sharma et al., 2020). Polyclonal human MAIT cells were generated in our lab from the flow cytometry-sorted and further expanded Vα7.2^+^CD161^+^CD4^−^CD8^+^ MAIT cells from the blood of healthy donors using the similar T cell expansion protocol (Huang et al., 2011; Sharma et al., 2020). The expanded polyclonal MAIT cells were saved as multiple aliquots at a cryogenic storage condition, and each aliquot was used for a one-time functional assay. In detail, blood samples from de-identified healthy donors without detectable infectious or non-infectious diseases were obtained with written informed consent at the Hoxworth Blood Center University of Cincinnati. We processed these human blood samples according to the protocols approved by the Institutional Review Board of the University of Cincinnati. We isolated human peripheral blood mononuclear cells (PBMCs) using the Ficoll-paque gradient (GE Healthcare) and removed the adhered monocytes after induced plastic adherence with cell culture flasks. The non-adhered PBMCs were washed twice with staining buffer (PBS with 2% FBS) and then blocked with anti-human Fc receptor antibodies, including anti-CD64 (mouse IgG1, clone 10.1), CD32 (mouse IgG2b, clone FUN-2), CD16 (mouse IgG1, clone 3G8), and additional Fc receptor blocking solution human TruStain FcX. PBMCs were further incubated with the combination of monoclonal antibodies, including phycoerythrin (PE)-Vα7.2, fluorescein isothiocyanate (FITC)-CD4 (OKT4), brilliant violet 711-CD8α (RPA-T8), and Allophycocyanin/Cyanine7 (APC/Cy7)-CD161 (HP-3G10), and sorted with flow cytometer FACSAria II (BD biosciences) for Vα7.2^+^CD161^+^CD4^−^CD8^+^ MAIT cells. The sorted CD8^+^ MAIT cells were expanded using feeder cells of irradiated human PBMCs (5, 000 rads), EBV-transformed B lymphoblastoid cell lines (7, 500 rads), together with stimulatory anti-CD3 antibody (clone OKT-3 at 30 ng/ml) and recombinant human IL-2 (1~2 nM). The expanded MAIT cells were harvested and confirmed with Enzyme-linked Immunospot assays.

### Enzyme-linked Immunospot (ELISPOT) assays of MAIT cell activation using bacterial eluates

As we reported (Sharma et al., 2017), ELSIPOT assays were performed using a MultiScreen sterile 96-well filter plate with 0.45 um pore size of hydrophobic polyvinylidene difluoride (PVDF) membrane (Millipore S2EM004M99), antigen-presenting cells K562.hMR1, and monoclonal or polyclonal MAIT cells. Briefly, the PVDF membrane was activated using 35% ethanol solution for 3 mins, washed with sterile H2O and PBS, then coated with purified anti-human interferon γ (IFNγ) antibody (Mabtech) for 16 hrs at 4°C. K562.hMR1 cells as antigen-presenting cells and monoclonal or polyclonal MAIT cells as responder T cells were mixed at a ratio of 2 × 10^4^ to 6 × 10^3^ cells in one well. Bacterial eluates (<3 kDa) were added at 3~5 ul/well, which can be normalized to 60~100ul bacteria culture. Upon co-culture for 16 hrs, cells were lysed and washed. The IFNγ^+^ MAIT cell spots were then developed with an indirect immunostain approach sequentially using a biotinylated anti-human IFNγ antibody (Mabtech), ExtraAvidin conjugated by alkaline phosphatase (Sigma), and substrates 5-bromo-4-chloro-3’-indolyphosphate/nitro-blue tetrazolium (BCIP/NBT, Sigma). We used CTL-ImmunoSpot S6 Micro Analyzer to visualize and quantify IFNγ^+^ MAIT cell spots. The blockade of responses was performed with an anti-MR1 antibody (clone 26.5, mouse IgG2a, at 2ug/ml) that blocks MR1-dependent MAIT cell activation (Huang et al., 2005; Huang et al., 2008; Huang et al., 2009). Anti-HLAI antibody (clone W6/32, mouse IgG2a, Biolegend, at 2ug/ml) was used as an isotype control for the anti-MR1 antibody (Sharma et al., 2020) and an irrelevant molecule control to block the effect of MHC class I proteins with structural similarity to MR1 (Wang et al., 2011).

### High-pressure liquid chromatography (HPLC) of bacterial eluates

The Shimadzu prominence HPLC system, containing a CBM-20A communication bus module, a DGU-20A3 degasser, a LC-20AD liquid chromatograph, a SPD-20A UV/VIS detector, and a FRC-10A fraction collector, was used for fractionation. Bacterial eluates (<3 kDa) for HPLC fractionation were injected at 75~150ul, which was normalized to 1.5-3 ml bacterial culture with OD_600_ absorbance of 1. Samples were separated using a diamond hydride column with TYPE-C silica (pore size 100 Å, particle size 4 um, length 150 mm, and diameter 4.6 mm, Cogent No. 70000-15P) linked to a guard column (pore size 100Å, particle size 4 um, length 20 mm, and diameter 4 mm, Cogent No. 70000-GC6). Mobile phase A contains 80% Optima H2O and 20% acetonitrile, and mobile phase B contains 100% acetonitrile. An aqueous normal phase gradient started with 95% of mobile phase B at 0-2 mins, changed to 80% at 4 mins, 70% at 21.5 mins, 10% at 22.5-23 mins, followed by column regeneration with mobile phase B to 95% at 25 mins. The flow rate was set at 500 ul/min to accommodate column size and allowed an easier collection and downstream preparation of samples. HPLC fractions were collected in an FRC-10A fraction collector using 4 ml glass tubes, and mobile phases were concentrated from 500 ul to around 50 ul using nitrogen gas evaporation. Upon evaporation, 5-10 ul of each HPLC fraction was used for functional confirmation using ELISPOT assays, and the remaining HPLC fractions were used for mass profiling analyses.

### Mass spectrometry-based comparative metabolomics of HPLC fractions

HPLC fractions of bacterial metabolite eluates were profiled using a Thermo Scientific™ Orbitrap Fusion™ Lumos™ Tribrid mass spectrometer controlled by XCalibur software. We applied the same diamond hydride column and the same aqueous normal phase gradient, which employed a nearly isocratic gradient with gradually increasing polarity, to further separate a single HPLC fraction. For each run, we injected 12 ul concentrated HPLC fractions, which were normalized to 0.5 ml of bacterial culture at an OD_600_ reading of 1. An additional additive chemical of 0.05% formic acid was applied in some mass profiling runs. Comparative mass profiling was set up using HPLC fractions from *M. tuberculosis* (H37Ra), *E.coli* (BL21), and *L. monocytogenes* (J1060), together with the solvent blank. Duplicate samples were injected for each HPLC fraction in the comparative mass spectrometry analyses. Reference compounds were run at a later confirmation stage using the same diamond hydride column and the same aqueous normal phase gradient. Ionization occurred with a heated-electrospray ionization source (H-ESI) with sheath gas, aux gas, and sweep gas of 35, 10, and 1 Arb, respectively. The Ion transfer tube temperature was maintained at 325°C and the vaporizer temperature at 325°C. The spray voltage was set at 3.5 kV for a positive ion mode. Spectral collection for MS1 was monitored between *m/z* 150-1000 with a normal ion trap scan rate using the detection type of Orbitrap at a resolution of 120K. The automatic gain control (AGC) target was set at 4 × 10^5^, microscans at 1, and radio frequency (RF) lens at 30% to collect profile data. Filter dynamic exclusion was applied for parental molecular ions with exclusion after 2 times, exclusion duration of 45 sec, a mass tolerance of 10 parts per million (ppm), and exclusion of isotopes. Some targeted collisional mass spectrometry analyses used Fourier Transformed Mass Spectrometry (FTMS, Thermo) under a similar HPLC condition and a matched collisional setting.

### Profile analyses of mass data

Mass profiles were initially visualized and manually compared using MZmine software and further converted into mgf and mzData files to be input for comparative metabolomic analyses of molecular ions and features associated with active fractions. Because Global Natural Product Social Molecular Networking (GNPS) was designed to analyze mass profiles of natural products (Olivon et al., 2017), we converted raw mass files to mgf files, input mgf files to the GNPS program, and used default settings to perform molecular network search (Wang et al., 2016). The molecular network was further input into the Cytoscape program to visualize the accurate mass of molecular ions as nodes, each of which annotated each molecular ion with an accurate mass and different bacterial sources. The cosine scores as edges measured the similarity of fragmentation patterns, with a cosine score of 1 representing identical spectra and 0 denoting no similarity (Nothias et al., 2020). Generally, a cosine score of 0.7 or higher measures similar fragmentation patterns with a minimal false positive alignment. To analyze the duplicate runs of each bacterium statistically, the updated XCMS program package (version 3.5.1) in the R platform performed peak picking with the centWave algorithm, quantification using the intensity-weighted mean algorithm, and retention time alignment by applying the script adapted from the published protocol (Huang et al., 2011). The mass profiles were aligned in the XCMS program across different bacteria and duplicate runs, using a signal-to-noise threshold of 10, *m/z* deviation of 5 ppm, frame width of mzdiff at 0.00001, a threshold count of 10000, a peak width of 10-60 s, and a bandwidth of 5. Results of XCMS analysis showed a list of molecular features with defined mass-to-charge *(m/z)* values, intensity counts of peaks, and aligned retention time. We then further annotated the molecular features associated with *M. tuberculosis* by defining the minimal intensities from *M. tuberculosis* (>10^5^ counts) and relatively low background intensities in the solvent blank (<10^6^ counts). The intensity of *M. tuberculosis-associated* molecular features was further defined with 50-fold higher intensity counts than those in solvent blank and 5-fold higher than those in *L. monocytogenes* or *E. coli*. Further, a comparative metabolome was shown using a volcano plot for molecular features obtained for *M. tuberculosis* and *E.coli* fractions with a significant p-value (<0.05) corrected by a false discovery rate (FDR). Thus, the integrated bioinformatics with mass profiling analyses provided comparative metabolomes and allowed us to associate individual molecular features with active HPLC fractions of *M. tuberculosis*.

### Structural prediction of bacterial metabolites using collision-induced dissociation mass spectrometry (CID-MS)

The molecular features associated with or uniquely presented in *M. tuberculosis* instead of *E. coli*, as shown in the volcano plot, were further examined using an extract ion chromatogram (EIC) of the targeted *m/z* values at a mass tolerance of 20 ppm. The molecular features that showed visible and significantly higher EIC from *M. tuberculosis* would be collided for structural prediction. For CID analyses at the MS2 level, we used isolation mode of quadrupole, isolation window at 2, activation type of CID, collision energy of 35%, activation time of 30 ms, activation Q at 0.25, detector type of ion trap using a normal scan rate, automatic gain control target at 1 × 10^4^, and microscans at 1 to obtain collisional centroid data. Candidate structures for parental ions were first predicted based on a match of *m/z* values between collided and calculated fragments. Reference 1-methylinosine (M=282.096, Medchemexpress Cat# HY-113139 from Fisher Scientific) was directly dissolved with Optima H2O at 10 ug/ul. This original stock was further diluted using Optima H2O to 2.5 ng/ul, and 20 ul dilution with 50 ng of 1-methylinosine was injected for mass spectrometry.

### Preparation of synthetic reference compounds for MAIT cell activation

Similar to the *M. tuberculosis* eluates (<3 kDa) potentially containing unstable active compounds, synthetic compounds could be unstable for cell activation. Guanosine was initially dissolved with DMSO at 50 ug/ul or 200 mM. Liposomes (Lipofectamine 3000, Invitrogen) were also applied at 1:100 dilution equally in DMSO and different guanosine concentrations. The 1-methyl-inosine was directly dissolved with Optima H2O at 10 ug/ul or with PBS at 40 mM. The 2-deoxy-5-formyluridine (M=256.070, Cat# ND29039 from Biosynth Carbosynth) was initially dissolved in DMSO at 25.6 ug/ul or 100 mM. Acetyl-6-formylpterin (M=233.055, Cat# 23303 from Cayman Chemical) was initially dissolved in ethanol at 1 ug/ul or with an additional culture medium to 1 mM (for example, 100ug with 100ul ethanol and an additional 329ul culture medium). Inosine (M=268.081, ACROS Organics Cat# EA AC122250250 from Fisher Scientific) or 9-(b-D-arabinofuranosyl)-hypoxanthine (M=268.081, Cat# NA03401 from Biosynth Carbosynth) were dissolved with Optima H2O to 100 mM. N2,N2-dimethyl-guanosine (M=311.123, Cat# ND05647 from Biosynth Carbosynth) was initially dissolved in DMSO at 40 mM. These initially dissolved stocks were further diluted with culture medium serially for cell assays. Chemical compounds were dissolved, diluted, and used freshly for cell assays by avoiding strong light and keeping at 4°C after dissolved in solution.

## Acknowledgement

We thank D. Branch Moody for suggestions on experimental design, data presentation, and writing; Shuangmin Zhang for conditioning ELISPOT assays; David Lewinsohn for MAIT cell clone D466F5; human blood donors of the Hoxworth Blood Center; and Robert Giulitto and Michelle Bailey for coordinating human blood samples. This work was supported by National Institute of Allergy and Infectious Diseases (AI115358, S.H.) and American Lung Association (IA-629987, S.H.).

## Author Contributions

Conceptualization: S.H. and T.H.; Investigation and assays: S.H., M.S., L.S., C.L., and Z.K.; methodology and resources: S.H., D.C., D.N., M.H., and S.C.; data curation: S.H. and S.C.; writing of original drafts: S.H.; draft editing: D.C., D.N., M.H., T.H., and S.C.; draft review: all authors; supervision, project administration, and funding acquisition: S.H.

## Declaration of interests

The authors declare no competing interests.

---

**Fig. S1.**
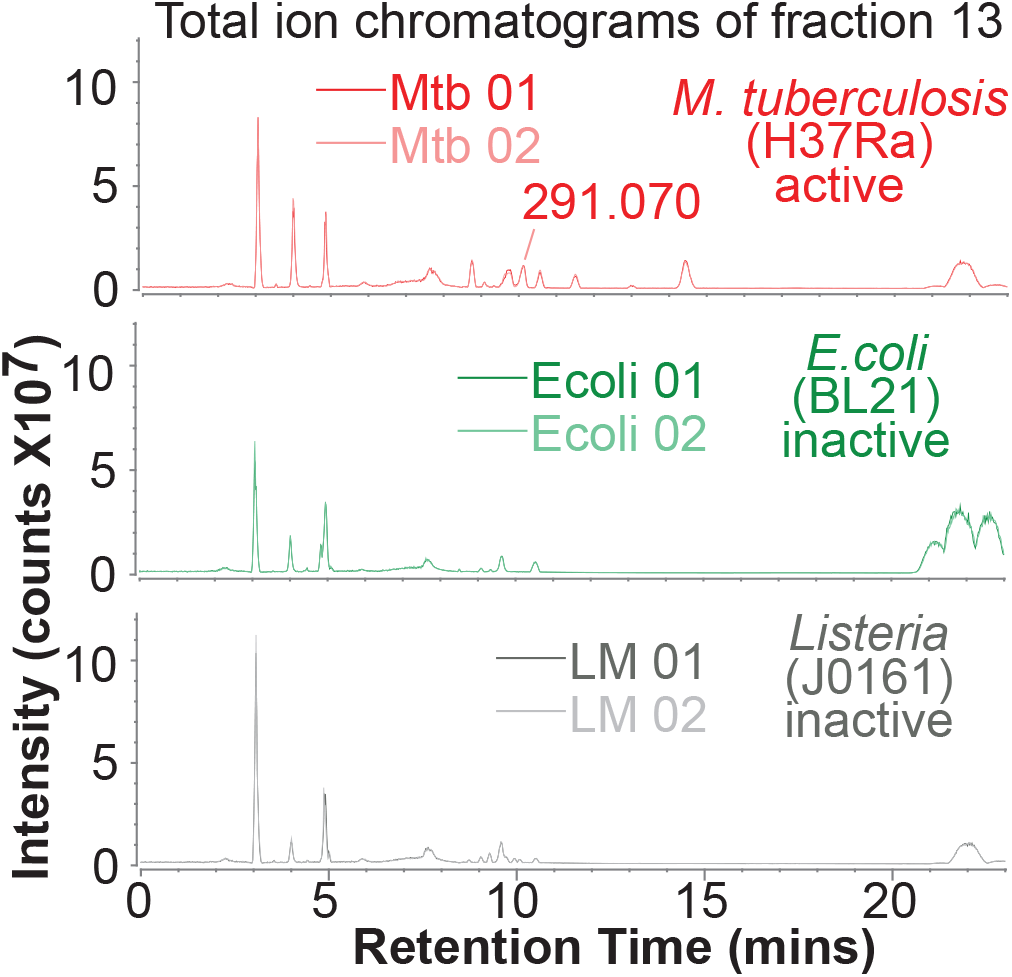
Total ion chromatograms of HPLC fractions 13. Duplicate runs were performed for each bacterium. An example base peak of *m/z* 291.070 enriches in *M. tuberculosis* fraction.

**Fig. S2.**
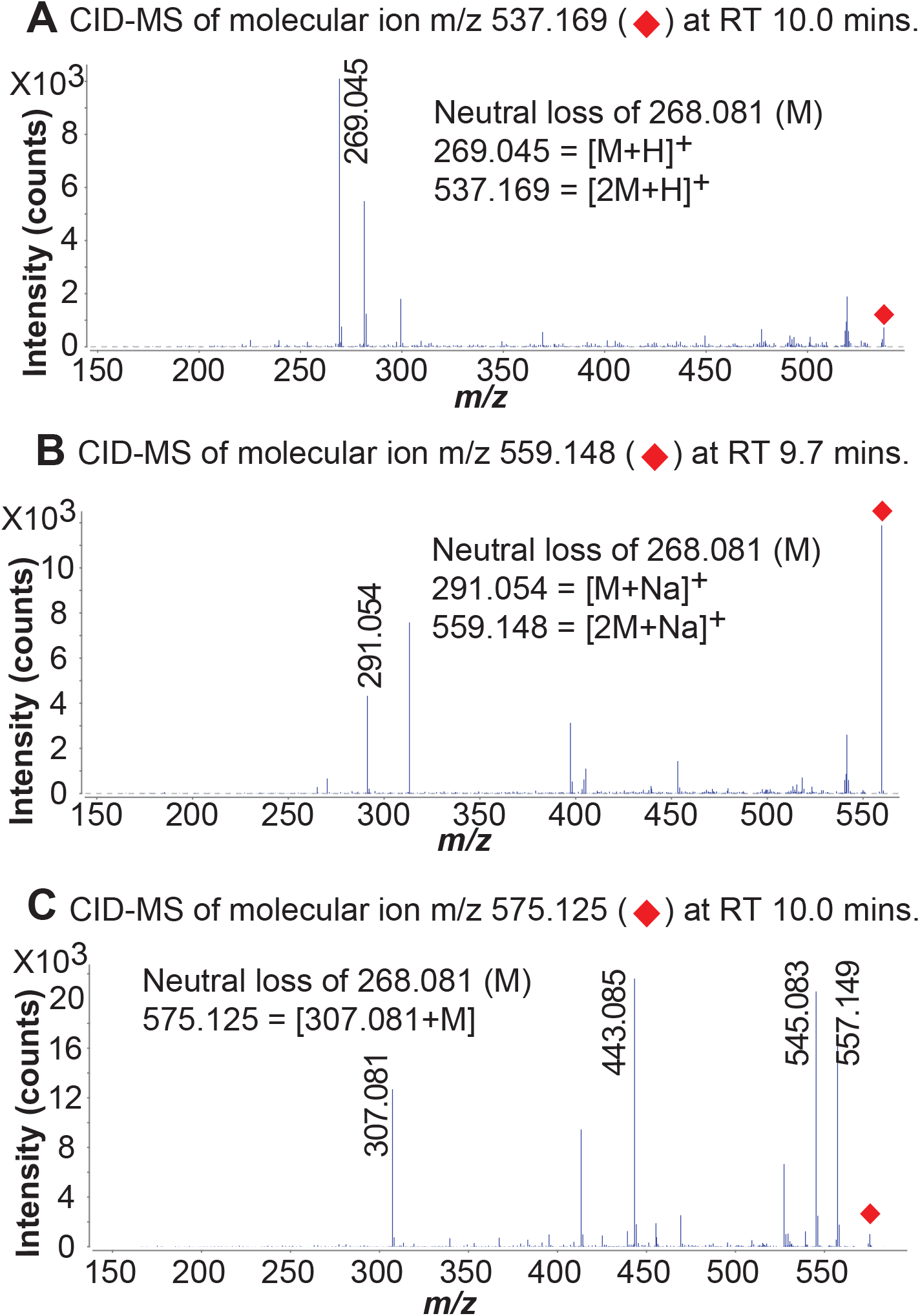
Molecular features with a common moiety. Enriched molecular features *m/z* 537.169 **(A)**, m/z 559.148 **(B)**, and *m/z* 575.125 **(C)** in *M. tuberculosis* share common neutral loss of mass 268.081.

**Fig. S3.**
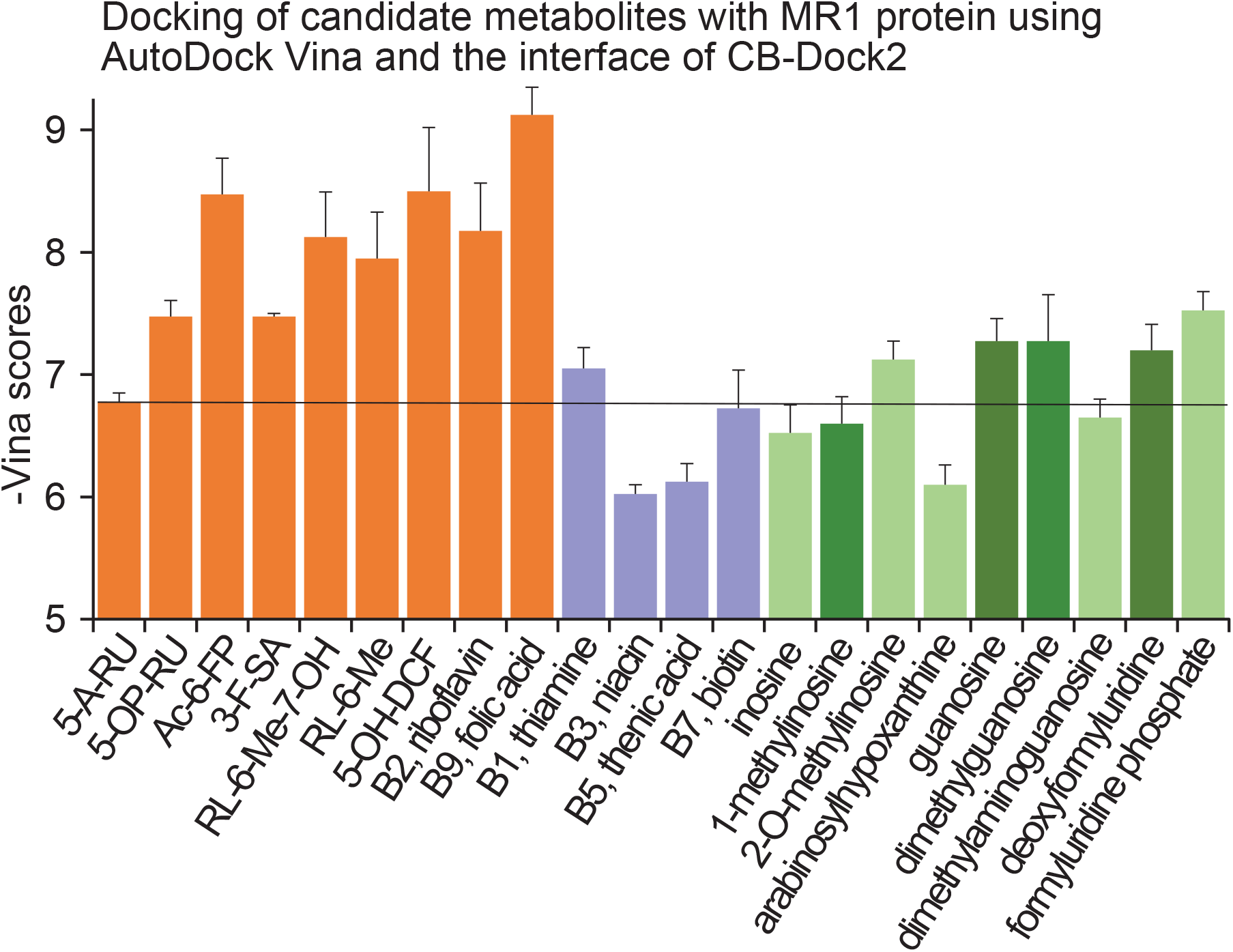
Autodocking of candidate nucleoside metabolites with four representative MR1 crystal structures. (4pj7 covalently bound to 5-OP-RU, 4l4v noncovalently bound to ribityllumazine, RL-6-Me-7-OH, 4nqd noncovalently bound to 5-OP-RU, and 4pjg covalently abound to Ac-6-FP). We used CB-Dock2 interface (https://cadd.lab-share.cn/cb-dock2/php/index.php) based on the Autodock Vina program (https://vi-na.scripps.edu/) to analyze metabolite docking in MR1 protein. The mean absolute values of Vina scores are plotted for each labeled metabolite upon docking with four noted MR1 crystal structures. Organge columns represent the known MR1 ligands or analogs. Blue columns represent negative controls from the vitamine B family. Green columns represent nucleosides either predicted from *M. tuberculosis* (light green) or as synthetic conserved analogs availalbe for functional assays. Vertical line represents a threshold that indicates the candidate metabolites potentially bind to the MR1 protein. This threshold was set upon the comparison of the Vina scores with the known MR1 ligands and negative controls. 5-A-RU: 5-amino-6-D-ribitylaminouracil, 5-OP-RU: 5-(2-oxopropylideneamino)-6-D-ribitylaminouracil, Ac-6-FP: acetyl-6-formylpterin, 3-F-SA: 3-formylsalicylic acid, RL-6-Me-7-OH: 7-hydroxy-6-methyl-8-d-ribityllumazine, RL-6-Me: 6-methyl-8-d-ribityllumazine, 5-OH-DCF: 5-hydroxy diclofenac.

## Notes

### Competing Interest Statement

The authors have declared no competing interest.

